# Fetal daily rhythms develop during pregnancy and entrain to the maternal circadian system

**DOI:** 10.1101/2023.06.30.547166

**Authors:** Keenan Bates, Elizabeth Sapiro, Jacob L. Amme, Ronald McCarthy, K.L. Nikhil, Sarah Speck, Varun Vasireddy, Ethan Roberts, Carmel Martin-Fairey, Miguel-E. Domínguez-Romero, Sandra Paola Garcia Cardenas, Sarah K. England, Erik D. Herzog

## Abstract

Circadian rhythms in gene expression and hormones are ubiquitous across species and differentiated cell types. This study aimed to determine when daily rhythms begin in the fetus and synchronize to the mother. We developed methods to monitor the expression of fetal PERIOD2 (PER2), a core circadian clock protein, in mice longitudinally from embryonic day (E)8.5 to E17.5 through *in utero* bioluminescence imaging. We found that embryonic PER2 expression increased rapidly throughout pregnancy and exhibited day-night rhythms from the start of our recordings at E8.5. The daily peak time of PER2 varied between pregnancies until it reliably peaked at night and synchronized to the mother starting around E15.5. Loss of fetal circadian rhythms associated with pregnancies that ultimately failed. Because maternal glucocorticoids have been implicated in fetal development and synchronizing circadian tissues, we tested their sufficiency to shift fetal daily rhythms in utero. Daily glucocorticoids injections over five days of late pregnancy advanced fetal PER2 rhythms *in utero* and blocking glucocorticoid signaling *in vitro* reduced PER2 synchrony between the maternal and fetal placenta by ∼40%. We conclude that fetal daily rhythms arise early in pregnancy and then synchronize with the maternal rhythm prior to birth depending, in part, on glucocorticoid signaling.

## Introduction

Circadian rhythms in gene expression and hormones generate daily cycles in behavior and physiology in many species and cell types [1]. In mammals, individual cells utilize transcription-translation feedback loops of core clock genes, including *Period2* (*Per2)*, to generate and maintain circadian rhythms [2]. In the adult, the suprachiasmatic nucleus (SCN) in the hypothalamus generates near-24 h rhythms, entrains to environmental cues such as the daily light-dark cycle, and regulates daily rhythms throughout the body [3]. When and where these daily molecular oscillations first arise in development is not known.

Daily rhythms in clock genes are not found in embryonic stem cells or induced pluripotent stem cells until differentiation [4–6]. Dedifferentiation back into the pluripotent stem cell state causes these rhythms to be lost [6]. *Ex vivo* studies have found circadian rhythms in a variety of fetal tissues [7–10]. For example, explants of fetal mouse liver and kidneys exhibit daily rhythms as early as embryonic day (E)13.5 (mouse gestation lasts 19-20 days) when the SCN does not yet exhibit rhythms. A day later, ∼10% of cells within the fetal mouse SCN develop circadian rhythms and, by E15.5, almost all SCN cells show synchronized daily rhythms [8–10]. However, *in vivo* data are not fully concordant. Early studies of fetal rat SCN showed day-night differences in metabolic activity from E19 onward (rat gestation lasts 21-22 days) [11,12]. *In vivo* measurements of mRNA and protein expression of clock genes in the fetal rat SCN and liver and whole mouse embryos either failed to detect rhythms or only detected rhythms in clock genes one or two days prior to birth [7,13–15]. This discrepancy may reflect differences between *in vitro* and *in vivo* induction of circadian cycling. However, prior *in vivo* approaches required sacrificing many animals and pooling data from multiple fetuses and pregnancies. This method can detect when rhythms are synchronized across individuals but cannot assess when individual fetuses develop rhythms. One prior study assessed fetal clock gene expression in rats using a real time reporter of Period1 transcription and found evidence that fetal rhythms may develop *in* utero. However, they did not have the temporal resolution to determine when daily rhythms develop [16]. As a result of the inconsistent results between studies, when circadian rhythms initiate in the fetus remains unknown.

A functioning maternal circadian system has been shown to be important in the normal development of the fetus and its developing circadian rhythm [9,17–20]. For example, rodents that experience constant light exhibited disrupted maternal melatonin rhythms, reduced fetal weights, and placental abnormalities while repeated 6-hr phase shifts led to a reduction in successful pregnancies [17,18]. Shift work during gestation has been associated with increased risk of preterm birth, placental complications, and pediatric obesity [21–26].

Pregnancy offers an interesting scenario as maternal and fetal circadian oscillators likely interact as fetal circadian rhythms develop. Maternal humoral signals, such as melatonin or glucocorticoids, may act to entrain or drive daily rhythms in the fetus [27,28]. Any entraining signal must pass through the placenta to interact with the fetus. Importantly, *in vivo* and *in vitro* experiments demonstrate that the placenta also exhibits circadian rhythms in clock genes and enzymes [28–31]. For example, daily rhythms in placental 11β-hydroxysteroid dehydrogenase (11β-HSD) inactivate glucocorticoids during rest. This allows active glucocorticoids to cross the placenta during wake, generating a signal that could potentially entrain the fetus [28,32,33]. Importantly, some of the potential entraining signals function in other developmental processes, such as glucocorticoid regulation of fetal lung development [34]. In pregnancies at risk of preterm birth, antenatal glucocorticoid treatments are used to accelerate fetal lung development [34–36]. However, the time-of-day of treatment is rarely considered which could reduce efficacy. Studies in mice (daily injections from E11.5 onward) and humans (2 daily doses at 24-34 weeks of gestation) showed that glucocorticoid treatments during gestation given out of phase relative to the endogenous rhythm caused diminished resilience to stress for the offspring [37]. Therefore, more research needs to be done to understand how the timing of glucocorticoid administration affects the development of fetal circadian rhythms.

This study aimed to measure maternal-fetal coordination of clock gene expression during pregnancy. We hypothesized that fetal circadian rhythms arise before the fetal SCN is developed and circadian rhythms synchronize to the mother after fetal SCN formation. We monitored fetal mouse clock gene expression *in utero* and found that fetal daily rhythms arise by E9 and gradually synchronize to the mother by E16. *In vitro* recordings revealed parallel changes in the time of daily peak clock gene expression in the uterus and placenta. The placenta exhibited intrinsic circadian rhythms beginning as early as E9. Administration of daily glucocorticoids to dams increased maternal-fetal synchrony while glucocorticoid receptor antagonists in vitro disrupted synchrony within the placenta. These results support the hypothesis that fetal circadian rhythms develop during pregnancy and entrain to the mother prior to birth, in part, through glucocorticoid signaling.

## Materials and Methods

### Animals

All experiments used mice maintained on a C57BL/6JN background. We utilized transgenic mice expressing a fusion protein of PERIOD2 and LUCIFERASE (homozygous PER2::LUC)[38]. Mice were housed in a 12h:12h light-dark cycle (lights on at 7am, defined as Zeitgeber Time 0, ZT0) in the Danforth Animal Facility at Washington University in St. Louis. Animals received *ad libitum* access to food and water throughout the experiments. Timed pregnancies were generated by pairing one male with two females overnight. Vaginal plugs the following morning defined embryonic day 0.5 (E0.5) and E1 ended with lights off (7 PM). All procedures were approved by the Animal Care and Use Committee of Washington University and followed National Institutes of Health guidelines.

### In utero imaging of fetal PER2 expression

We mated wild-type females to homozygous PER2::LUC transgenic males. We imaged dams twice daily within 1 hour of lights on and lights off from E8.5-E14.5 and every 4 h from E14.5-E17.5. We used a depilatory cream (Nair, Church & Dwight) to remove fur from the dam’s abdominal area on E7.5. Dams received D-luciferin either as an injection (150mg/kg body weight; Xenolight, Perkin Elmer) 10 min prior to imaging or in their drinking water (10 mM, Gold Biotechnology). We replaced drinking water every 3 days. Anesthetized (2% isoflurane vaporized in 1 L/min O_2_) dams were imaged (IVIS Lumina III, Perkin Elmer) with their ventral side facing the charge-coupled device camera (15 cm field of view, 8×8 binning, f stop 1, 10-min exposure from E8.5-11.5 and 2-min exposure from E12.5-E17.5). Total photon flux (photons/s) was measured from *in utero* bioluminescence using fixed regions of interest (ROIs) over the abdomen of dams in Living Image 2.6 (Perkin Elmer).

To compare fetal PER2 expression in mice that were not handled, we measured bioluminescence from a subset of mice housed individually in constant darkness beginning on E13.5 in a light tight box (Lumicycle In Vivo, Actimetrics) containing two photomultiplier tubes (Hamamatsu H8259-01) [39]. Dark counts were collected for 1 min every 15 min by closing the programmable shutter for 1 min. We checked animals daily to ensure that they had appropriate food and water. We summed bioluminescence counts per minute from the two sensors following dark count subtraction.

To assess whether fetal PER2 rhythms within a pregnancy were synchronized, we took the bioluminescence images collected every 4 hours (14.5-E17.5) and aligned them in ImageJ software (NIH). These images were converted into a video for each pregnancy. Three pregnancies (1 control and 2 corticosterone injected) were excluded from this analysis because of a small litter size (1-2 pups). The videos underwent pixel-based analysis using a custom Python script to measure instantaneous phase and the synchronization index. We measured intensities by implementing a multidimensional gaussian filter in SciPy tools using established methods to generate a time series of pixel intensities [40]. We used pyBoat to perform wavelet analysis and measure instantaneous phase and sync index [41]. Code for data processing is available on request.

### In vivo corticosterone treatment

A subset of mice received 20 mg/mL corticosterone (50 mg/kg, Sigma-Aldrich) or vehicle (polyethylene glycol 400, Sigma-Aldrich) daily at 7:00am from E13.5-E17.5.

### *In Vitro* bioluminescent recording

Pregnant and non-pregnant PER2::LUC mice were killed with CO_2_, followed by cervical dislocation, at E9, E12, E15, E18, metestrus, and diestrus between ZT2 and ZT6. For estrous staging, we collected vaginal smears on glass slides and observed cell types using bright field microscopy 2 hours prior to tissue collection. At each time point, the placenta (if applicable), uterus, and cervix were extracted and placed in chilled Hank’s buffered salt solution (HBSS, Sigma). The placentas were sliced longitudinally and cut into cross-section explants containing all the layers of the placenta (decidua, junctional zone, and labyrinth zone) or further dissected into the individual layers of the placenta (∼4mm^3^). Uterine and cervical samples were cleared of fat and cut longitudinally into ∼2 x 2 mm pieces. All explants were placed on 0.4 mm MilliCell membrane inserts (Millipore) in 35 mm Petri dishes (BD Biosciences) containing 1 ml of DMEM (Sigma-Aldrich, pH 7.2), supplemented with 25 U/ml penicillin, 25 g/ml streptomycin (Invitrogen), 10 mM HEPES (SigmaAldrich), 2% B27 (Invitrogen), 0.35 g/L NaHCO3 (Sigma-Aldrich), and 0.1 mM beetle luciferin (BioSynth) and sealed with vacuum grease.

Placental, uterine, and cervical explants were transferred to light-tight incubators at 36^◦^C and placed under photomultiplier tubes (HC135-11MOD; Hamamatsu). Bioluminescence was recorded in 10-minute bins for 5 days. Bioluminescent traces were detrended by subtraction of the 24-hour running average and days 2-5 were fitted with a damped sine function (Chronostar version 1.0). Traces with a period between 18-30 hours and a coefficient of correlation >0.70 were considered circadian. The period was calculated using Chronostar software based on days 2-5 of the recording. Amplitude was determined using a MatLab code analyzing raw data from 36-60 hours after the start of the recording. Phase was determined based on the average time of peak PER2 expression for days 2-5 of the recording using MatLab. All phase data are reported in Zeitgeber Time (ZT) with ZT0 corresponding to lights on for the day of surgery. Phases were analyzed using Oriana (version 4.02). Average phase was calculated by vector addition. A probability of 0.05 from the Rayleigh test was used to indicate whether samples were significantly clustered.

To analyze whether glucocorticoids affected circadian rhythms in the placenta, a subset of E12 placental cross-sections were placed in a light-tight incubator at 36^◦^C under a CCD camera (Onyx, Stanford Photonics). These placentas were immediately cultured with either a synthetic glucocorticoid, dexamethasone (2 µL of 50 µM dexamethasone in 1 mL of recording media, 100 nM, Sigma-Aldirch), a glucocorticoid receptor and progesterone receptor antagonist, mifepristone (1 µL of 1 mM mifepristone dissolved in ethanol, 1 µM, Sigma-Aldrich), vehicle (0.01% ethanol in ddH_2_O), or a 30-minute pretreatment of 1 µM mifepristone before adding 100 Nm dexamethasone to the recording media. Images were collected every 30 minutes for 5 days and 2 images were averaged using ImageJ software (NIH) to generate one image per hour. Bright noise and cosmic radiation were filtered out using adjacent frame minimization in ImageJ.

The processed videos underwent pixel-based analysis using a custom Python script to calculate percent rhythmicity, measure instantaneous phase, and the synchronization index.We measured intensities by implementing a multidimensional gaussian filter in SciPy tools using established methods to generate a time series of pixel intensities [40]. We analyzed the time series for rhythmicity using MetaCycle [42]. Percentage of rhythmic pixels was calculated as the number of pixels reported by MetaCycle as significantly rhythmic (BH.Q < 0.01) with a period between 18h and 30 h / number of analyzed pixels with non-zero pixel intensities x 100. We used pyBoat to perform wavelet analysis and measure instantaneous phase and sync index [41]. Code for data processing is available on request.

### Statistical analysis

All results are presented as mean ± S.E.M. We used 2-way ANOVA to compare synchrony indices between corticosterone injected and control mice. We used Rayleigh’s Uniformity test for statistical analysis of the distributions of daily peak PER2 times from tissue explants. A p-value <0.05 corresponds to a significant clustering of peak PER2 times between samples. We used 1-way ANOVA to compare changes in period and amplitude at different gestational ages for explanted tissues. We used 1-way ANOVA with Tukey’s multiple comparsions test to compare sync indices between treatment groups.

## Results

### Fetal PER2 increases in level and day-night variation with gestation

To characterize the development of fetal clock gene expression *in utero*, we mated homozygous PER2::LUC males to wild-type female mice. As a first step, we imaged 10 pregnant dams under anesthesia following luciferin injection. We sacrificed one pregnant female following a luciferin injection on E17 and established that bioluminescence was derived exclusively from the fetus, placenta, and umbilical cord. When we imaged these pregnancies repeatedly, we found that 4 successfully gave birth (**Figures 1 and S1**), three resorbed the conceptus and three failed to initiate labor. Importantly, successful pregnancies had higher mean and day-night amplitude PER2 expression. We conclude that daily rhythms in fetal clock gene expression are associated with successful birth outcomes.

**Figure 1.**
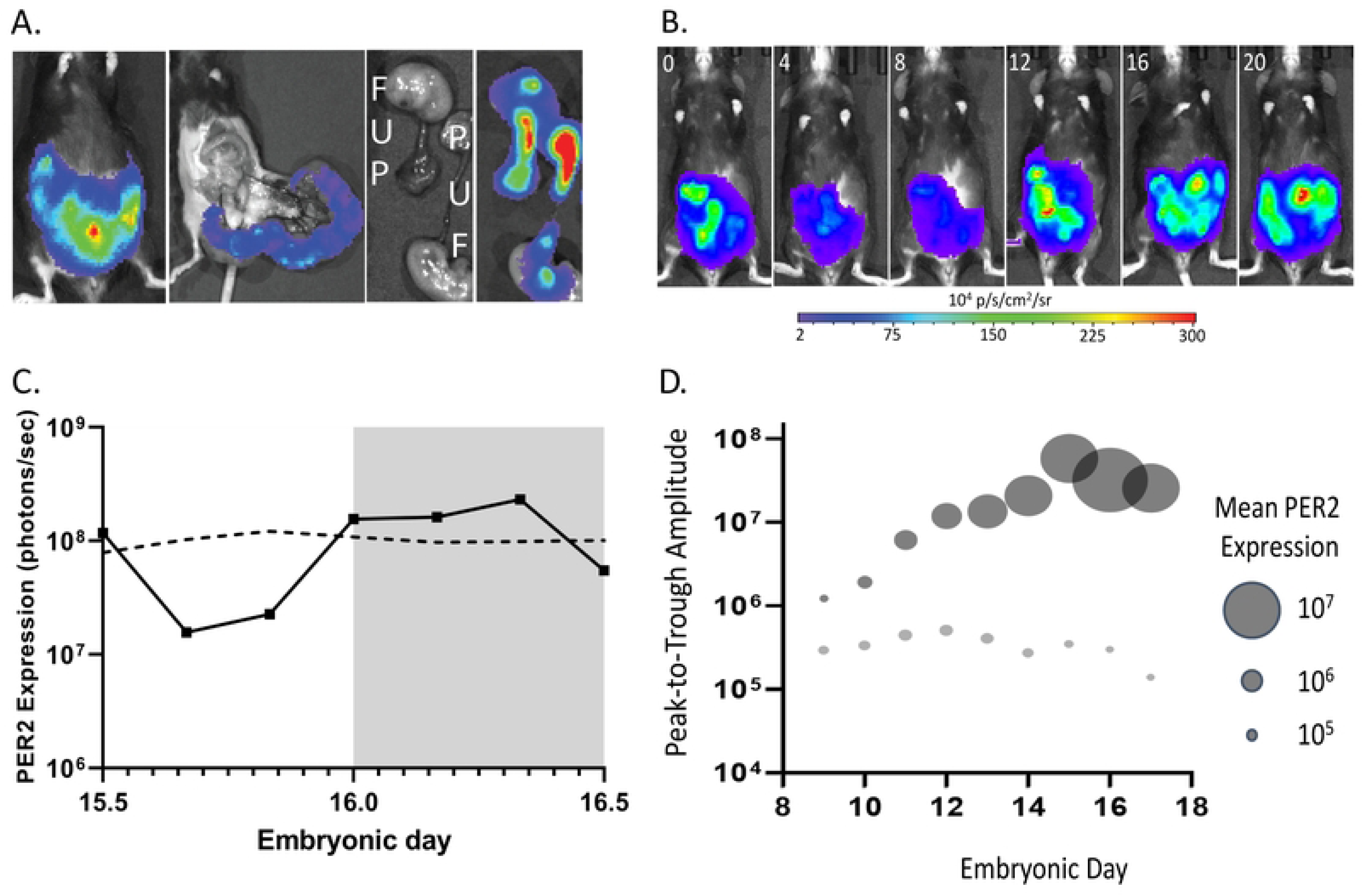
In utero imaging of fetal PER2 reveals that successful pregnancies associate with development of high amplitude circadian rhythms. A) Representative bioluminescence images of a E17.5 pregnant female following luciferin injection showing PER2 expression in utero, restricted to the uterus, and localized to two fetuses (F) and their placentae (P), and umbilical cords (U). B) Bioluminescence imaged every 4 h from a representative pregnant dam varied with time of day on E17. C) PER2 expression quantified as bioluminescence (solid line) around the running mean (dashed line) from the pregnancy in B. D) Through gestation, daily amplitude and mean (circle diameter) fetal PER2 expression increased in successful pregnancies (grey circles, n=4). Failed pregnancies did not show the increased daily mean and peak-to-trough amplitude of fetal PER2 (open circles, n=6).

### Fetal PER2 expression develops day-night differences prior to SCN maturation and synchronizes to the mother before birth

With a 60% pregnancy failure rate using repeated luciferin injections, we tested the efficacy of substrate delivery in the drinking water based on prior reports [39,43]. Similar to those reports, we found drinking rhythms caused negligible changes in day-night rhythms of PER2 bioluminescence (**Figure S2**). We, therefore, provided wild-type females (n=7) pregnant with PER2^Luc/+^ pups continuous access to luciferin-supplemented water starting the day prior to imaging. We imaged dams every 12 h from approximately E8.5 to E14.5 and then every 4 h until E17.5 (**Figure 2** and **Figure S3**) or every second from E14.5 to E19.5 (**Figure S2**). We found fetal PER2 expression increased exponentially during pregnancy with day-night differences starting between E8.5 and E9.5 in all pregnancies, several days before neurogenesis in the embryonic SCN [44].

**Figure 2.**
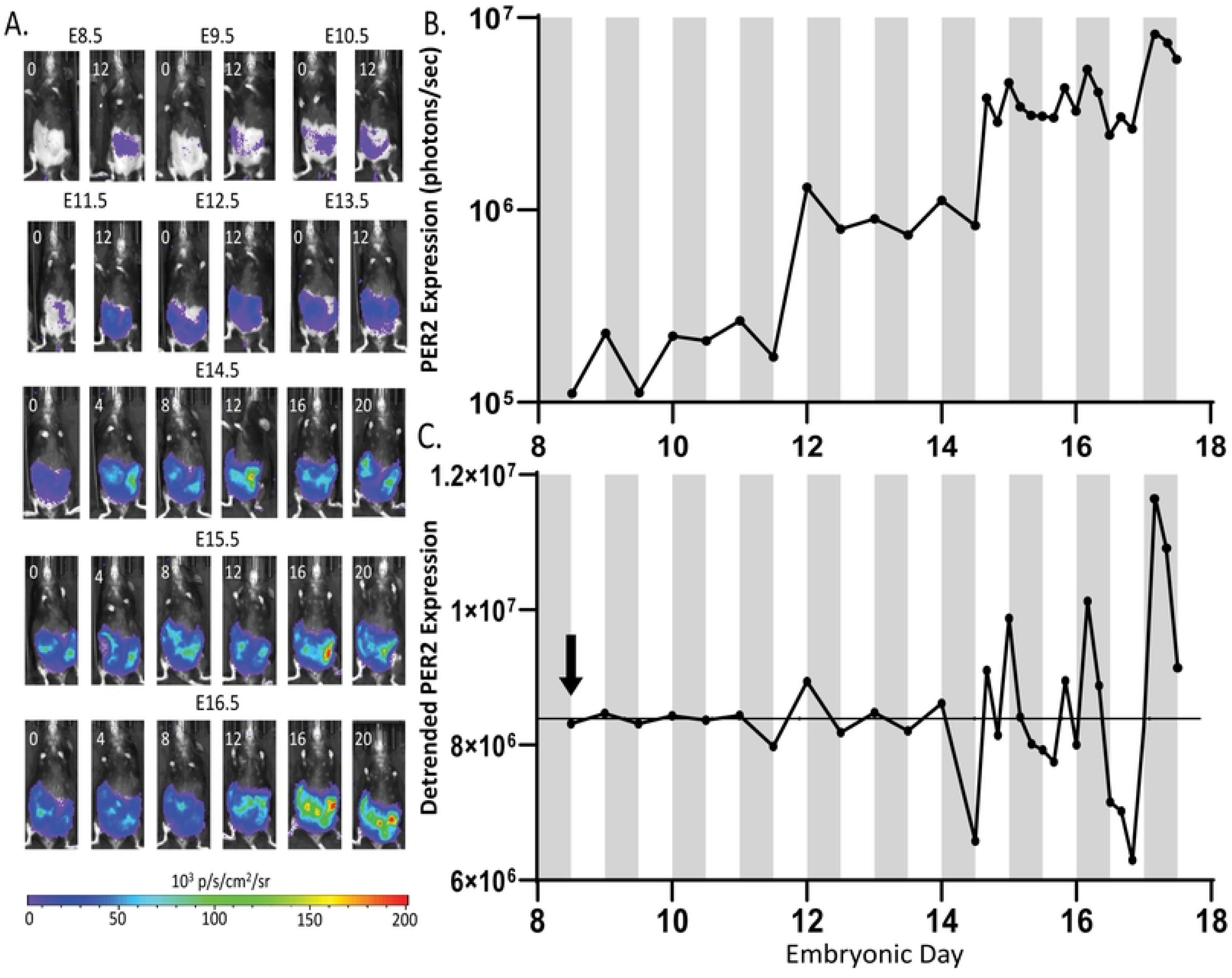
Day-night differences in fetal PER2 expression begins as early as E8.5. A) Sequential images from a representative dam show fetal PER2 expression was detectable on E8, varied from dawn (ZT0) to dusk (ZT12) and increased throughout pregnancy. B) Fetal bioluminescence increased exponentially in this representative dam. C) After subtracting the exponential increase from the data, day-night differences in PER2 expression were apparent starting on E8 (arrow) presented as a time series. Note that fetal PER2 expression gradually synchronized to peak at night during the last days of pregnancy.

We next examined whether fetal rhythms synchronize to the mother. First, we averaged PER2 expression from the individual pregnancies and found that, beginning on E15.5, lower expression reliably occurred during the day compared to the night (**Figure 3**). We compared the time of daily peak expression in each pregnancy and saw a clear increase in synchrony, as measured by the Rayleigh statistic, as gestation progressed. Recordings from two freely moving wildtype females pregnant with heterozygous PER2::LUC fetuses similarly revealed fetal PER2 expression was circadian from E14 and reliably peaked around subjective dusk by E15.5 (**Figure S2**). These results suggest that the fetus develops rhythms early in pregnancy but doesn’t fully synchronize to the mother until E15.5.

**Figure 3.**
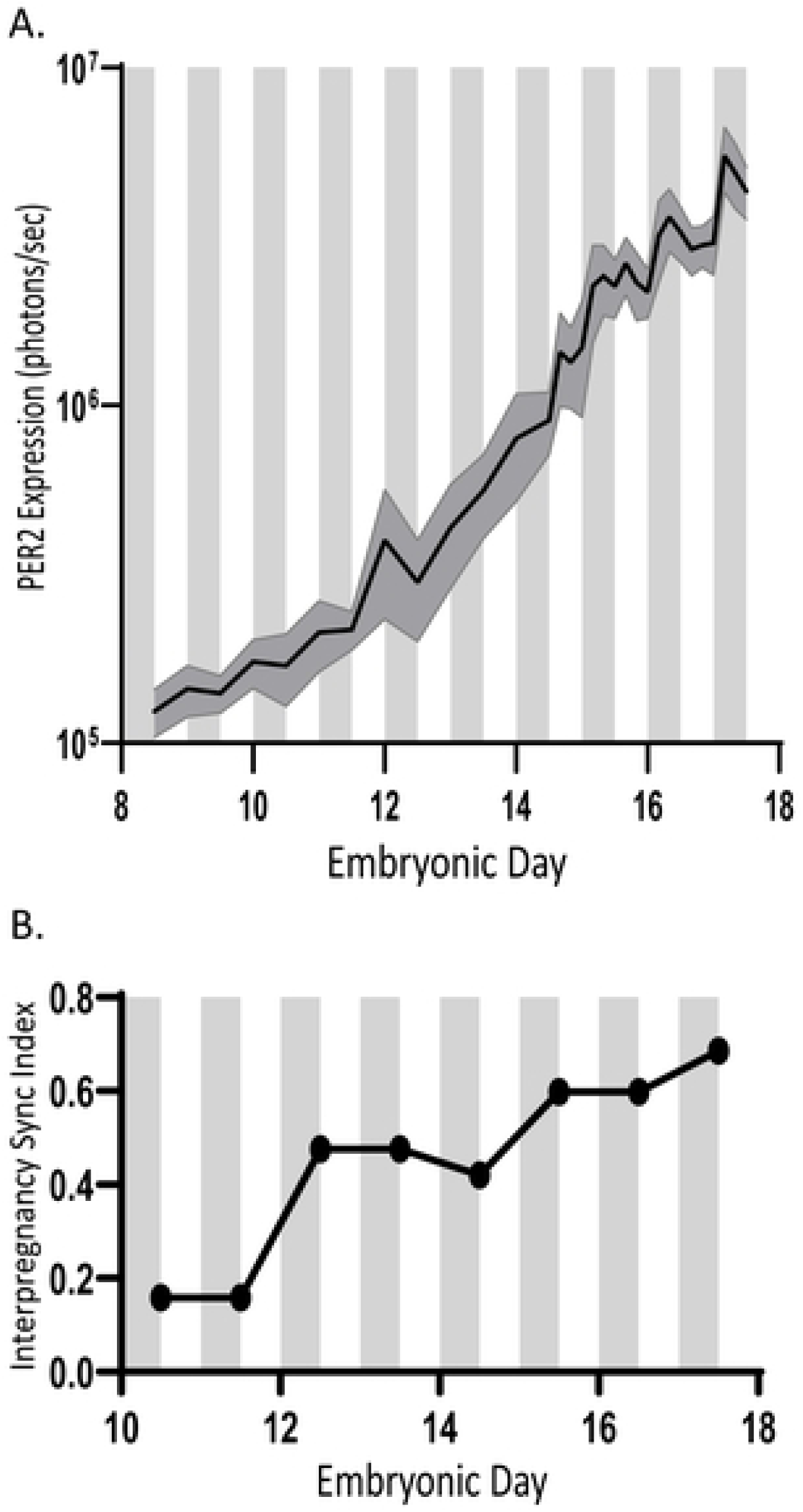
*In utero* PER2 expression exhibits day-night fluctuations in expression beginning at E8.5 and interpregnancy synchrony increases as pregnancy progresses. A) Average fetal PER2 expression from pregnancies imaged from E8.5 to E17.5 (n=6-7). Averaged results show day-night differences beginning early in the pregnancy, no coordinated rhythms in expression between E13-E14, and consolidated rhythms that peak at night from E15-E17. B) Interpregnancy synchrony index based on peak PER2 phase for each pregnancy (n=7) increased as pregnancy progressed. We measured synchrony starting at the earliest timepoint where all pregnancies exhibited rhythms (E10).

### Glucocorticoids entrain fetal circadian rhythms

Past research has shown that glucocorticoids can act as an entraining signal in the fetal SCN [28]. Because glucocorticoids typically peak around dusk in mice [45], we hypothesized that elevating corticosterone levels around dawn would shift the timing of fetal peak PER2 expression compared to controls. Following procedures from prior studies [37], we injected 50 mg/kg corticosterone each day for five days at ZT0 from E13.5-E17.5 in pregnant wildtype mice. We imaged the PER2::LUC fetuses every 4 h starting on E13.5 (n= 4) or E14.5 (n=3) until E17.5 (**Figure 4** **and Figure S4**). In corticosterone-injected mice, we found that the time of daily peak for fetal PER2 expression shifted by 4-8 h from E13.5 (first injection) to E17.5 (last injection) compared to control pregnancies (same mice as in Figure 2). Intriguingly, for three pregnancies, fetal PER2 expression was near its daily peak when the first corticosterone injection occurred, resulting in peak PER2 during early subjective night on E17. In contrast, the other four pregnancies had fetal PER2 near its daily minimum when the first corticosterone injection occurred, resulting a later time of peak PER2 on E17(**Figure S4**). To test whether corticosterone synchronized daily rhythms among fetuses within each pregnancy, we performed a continuous wavelet transform of bioluminescence in each pixel within the uterus of each dam (**Figure 4)**. Raster plots and intrapregnancy synchrony indices showed corticosterone increased synchrony of PER2 among fetuses within each pregnancy. Daily peak fetal PER2 expression rapidly aligned across pups within each dam during daily corticosterone injections compared to the slower synchronization of controls. Thus, glucocorticoid administration out-of-phase with endogenous daily rhythms accelerates synchrony among pups but also shifts the time of peak fetal PER2.

**Figure 4.**
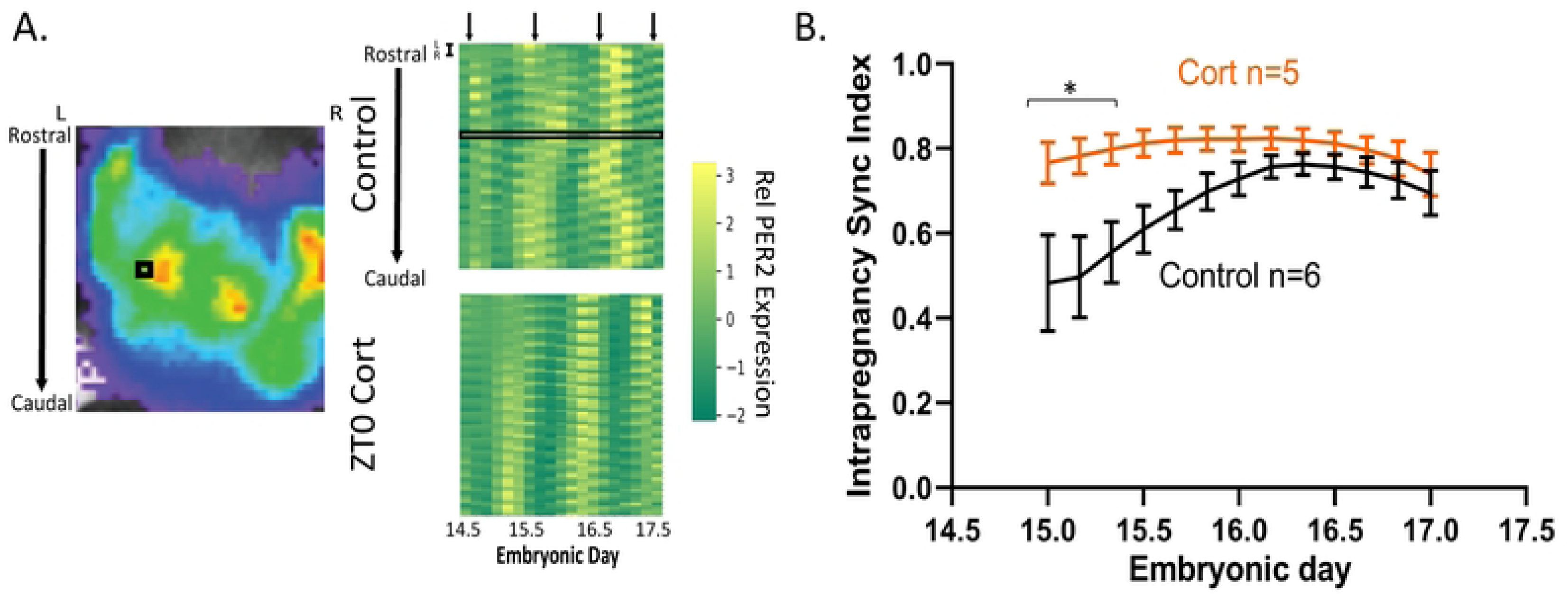
Daily corticosterone injections synchronize daily PER2 expression among *in utero* fetuses. A) Fetal PER2 expression within the uterus of a representative dam illustrating the location of one pixel near a fetus (black square). Raster plots show PER2 expression from E14.5-E17.5 for two representative pregnancies either untreated or injected with corticosterone at dawn. Note that the corticosterone-treated pregnancy shows increased clustering of daily peak PER2 expression by the end of the recording. The bioluminescence recorded from the representative pixel is highlighted in the black rectangle. B) Within pregnancy synchrony indices from control (n=6) and corticosterone-injected (n=5) dams show that corticosterone synchronized fetal PER2 faster than controls which synchronized as pregnancy progressed (2-way ANOVA; mean ± S.E.M).

### Circadian coordination between maternal and placental tissues

With strong evidence for synchronization of maternal-fetal clock gene expression in utero, we next tested for circadian communication between specific maternal and fetal tissues during pregnancy. We collected uterine, cervical, and placental explants at four embryonic ages (E9, E12, E15, and E18) and two estrous stages (metestrus and proestrus) and recorded their PER2 expression *in vitro*. All tissues exhibited intrinsic circadian rhythms at each of the stages of pregnancy (**Figure 5**). The daily amplitude of PER2 expression decreased with gestational age in the uterus (E9 vs E18, p= 0.008) and placenta (E9 vs. E15, p= 0.002 and E9 vs. E18, p= 0.0005; **Figure S5**). These results suggest that maternal and fetal tissues possess intrinsic circadian rhythms that could interact to affect pregnancy outcomes.

**Figure 5.**
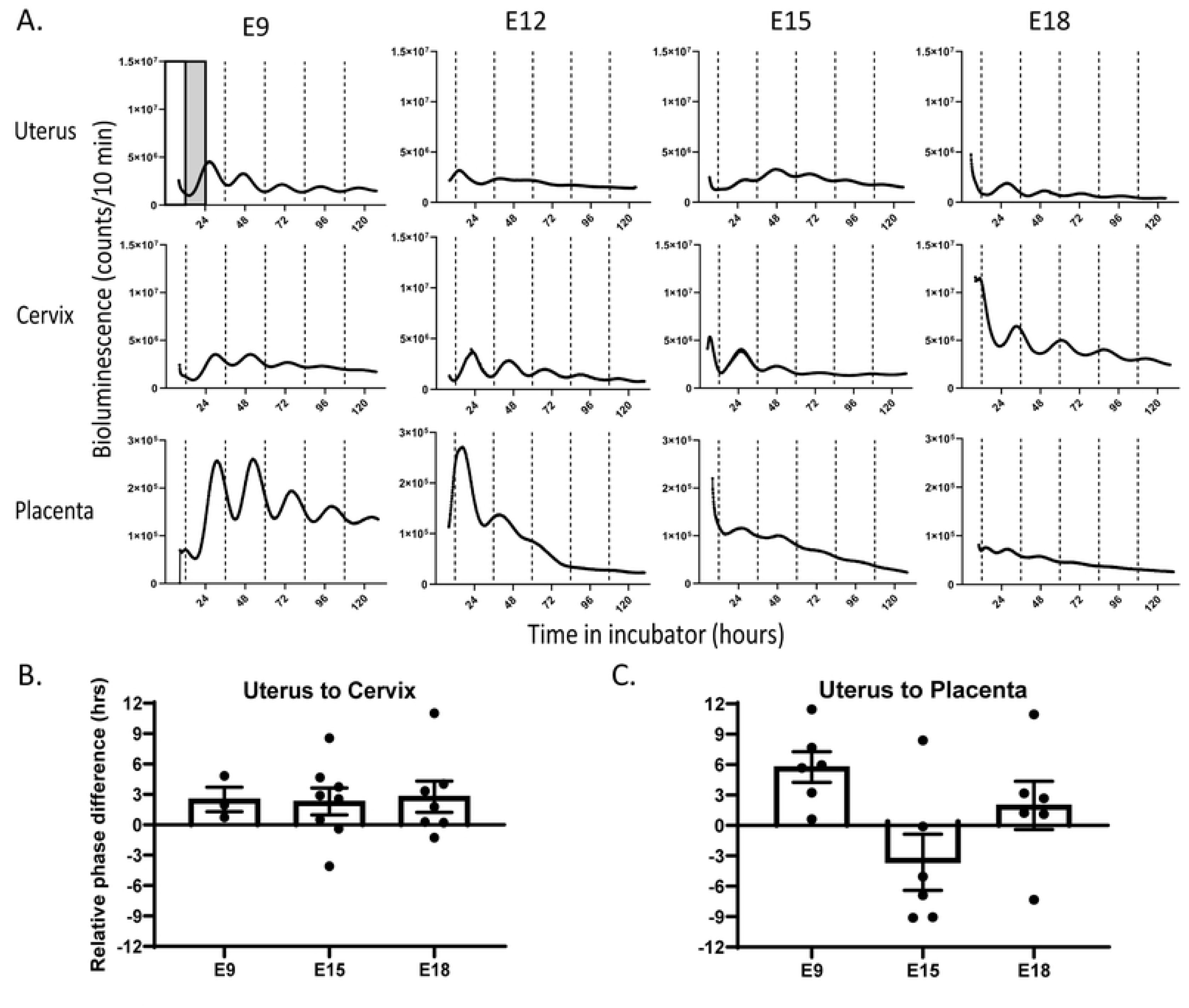
Coordinated daily rhythms intrinsic to the uterus, cervix, and placenta. A) Representative bioluminescence recordings from uterus, cervix, and placenta exhibited circadian rhythms in PER2 expression over 5 recording days when explanted at E9, E12, E15 or E18. Relative to the prior light-dark (white-grey bars) cycle, the time of peak PER2 expression depended on the tissue and embryonic stage. B) Within the same pregnancy the uterus peaked about 2.5 h before the cervix and C) had a phase relationship with the placenta that changed dramatically with gestational age.

We next examined the time of daily peak PER2 expression as a function of tissue type, estrous and pregnancy (**Figure S6**). PER2 expression in uterus peaked around mid-morning during metestrus and mid-afternoon during proestrus, similar to the findings from prior research [46]. The cervix showed a similar shift in time of peak PER2 from metestrus to proestrus. During pregnancy, the uterus consistently peaked about 2.5 h before the cervix from the same dam and between dams. In contrast, the placenta peaked after the uterus from the same pregnancy at E9 and E18, but their phase relationship was unreliable at E15. Intriguingly, at E15, the time of peak PER2 in the placenta and uterus was random when comparing between pregnancies. These results indicate that maternal and fetal tissues coordinately change their time of daily PER2 expression with pregnancy stage.

### Placental layers show intrinsic and coordinated circadian rhythms

To determine whether maternal or fetal layers contribute to circadian rhythms in the placenta, we explanted the placenta and compared PER2 expression in the intact placenta and isolated maternal decidua, fetal junctional zone, and fetal labyrinth zone (**Figure 6** and **Figure S7**). All three layers exhibited circadian rhythms when explanted at E12, 15 or 18. Imaging of the E12 placenta revealed a daily wave of PER2 expression from the maternal to fetal layers and higher amplitude circadian rhythms in the maternal placenta. All three layers also exhibited circadian rhythms when isolated from each other at E15 or 18 with the decidua reliably peaking in the early subjective night. Rhythms in the junctional and labyrinth zones were lower amplitude and less reliable in their times of peak expression. These data indicate that the decidua may coordinate circadian timing across the placenta.

**Figure 6.**
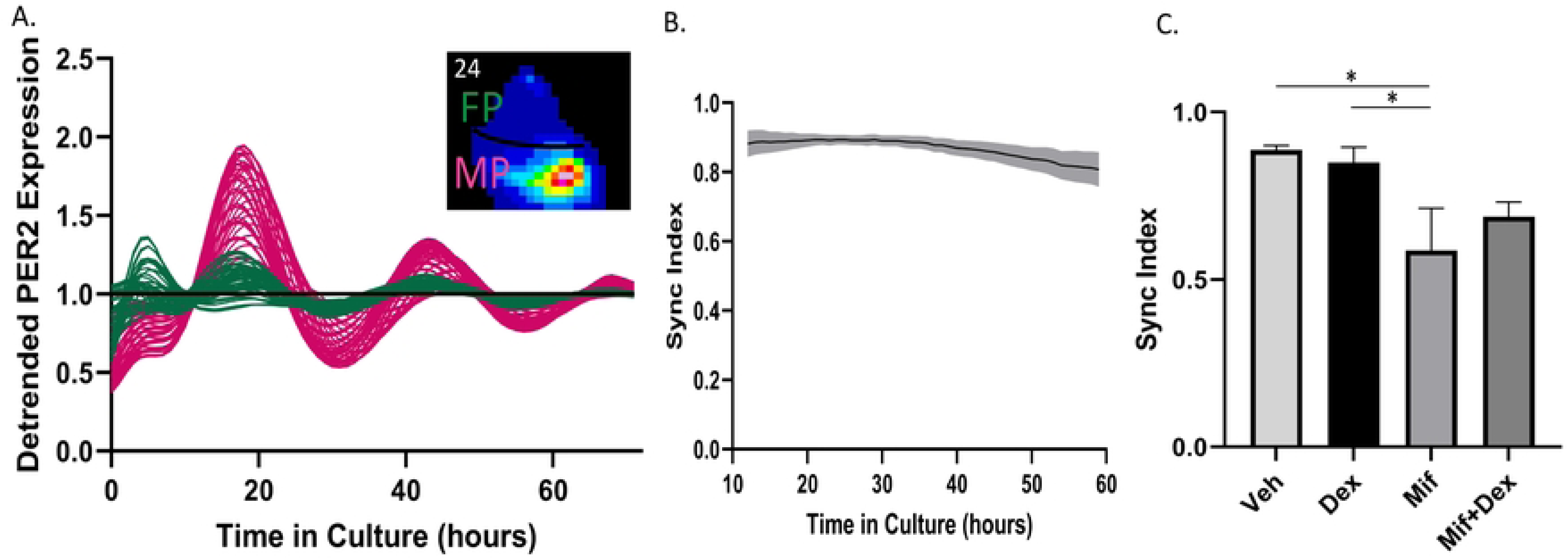
Synchronized circadian rhythms between maternal and fetal placenta depend on glucocorticoid signaling. A) Daily PER2 rhythms recorded over 5 days from 50 representative pixels within the E12 maternal and fetal placenta. Note the detrended bioluminscence from the maternal placenta showed higher amplitude rhythms. Inset shows PER2-driven bioluminescence from the representative maternal (MP) and fetal placental (FP) layers. B) Cells maintain high circadian synchrony throughout the placenta (n=5 vehicle-treated explants). C) A glucocorticoid antagonist (mifepristone, n=6) reduced synchrony between placental cells when compared to vehicle-and dexamethasone-treated placenta (n=5, p= 0.037; n=6, 0.049, respectively; One-way ANOVA with Tukey’s multiple comparsions; mean ± S.E.M).

To directly test for circadian communication between layers, we treated E12 placenta with a glucocorticoid agonist (100 nM dexamethasone), a progesterone/glucocorticoid antagonist (1 µM mifepristone), a combination of mifepristone and dexamethasone, or vehicle (0.01% ethanol). Mifepristone decreased synchrony among circadian cells within the placenta compared to vehicle or dexamethasone (**Figure 6C**, p < 0.05). We conclude that glucocorticoid signaling modulates synchrony between the maternal and fetal layers of the placenta.

## Discussion

Our results indicate that circadian rhythms develop at an early stage in embryonic mouse development. Using longitudinal imaging, we found day-night differences in PER2 expression begin by E8.5 and that gradually synchronize to the mother by E15. This is the earliest reported onset of daily rhythms in mammalian development [7–9,13]. *In vitro* recordings of the placenta showed intrinsic circadian rhythms at E9, the earliest stage we tested. *In vivo* administration of glucocorticoids enhanced synchrony of maternal-fetal daily rhythms while *in vitro* glucocorticoid antagonists disrupted circadian synchrony between maternal and fetal placenta. We posit that fetal circadian rhythms develop during pregnancy and then entrain to the mother prior to birth, in part, through maternal glucocorticoids.

Longitudinal imaging of pups in the same pregnancy has its advantages and disadvantages. We found that, in pregnancies that led to resorptions or failed to initiate labor, fetal PER2 expression levels and amplitudes were reduced compared to term pregnancies. The role of PER2 expression or circadian rhythms in pregnancy and development remains unclear. The stress from the repeated handling and luciferin injections may have put pregnancies at risk. By delivering luciferin in the drinking water, we increased our rate of successful deliveries from 60% to 100%, consistent with prior studies [39,43]. Prior *in vitro* research showed several fetal tissues expressed PER2 rhythmically as early as E13.5 but cannot rule out that rhythms were induced by surgical isolation [7–9]. Prior *in vivo* measures of fetal *Per2* mRNA, based on pooled data from multiple fetuses and pregnancies sacrificed at defined embryonic stages, found no rhythms in utero or evidence for weak daily rhythms one or two days before birth [7,13–15]. To detect rhythms, this approach requires synchronous daily rhythms between fetuses within and between pregnancies. We detected daily rhythms early in pregnancy that synchronized to the local light cycle around E15.5. Future experiments should be able to track individual fetuses and fetal cell types using conditional luciferase expression.

Maternal-fetal circadian communication appears in the early placenta and becomes synchronized to the light-dark cycle around E15.5. This timing corresponds with the rapid decline in HSDIIβ2 levels between E14.5 and E16.5 [47] which allows the high levels of maternal glucocorticoids [48] to escape inactivation by 11β-HSD2 [32]. We found that daily injections of glucocorticoids, which can saturate 11β-HSD2 and interact with the fetus [37], accelerated circadian synchrony among pups in utero. In mice and humans, dexamethasone injections given out of phase with the normal rise in glucocorticoids has been associated with increased anxiety-like behaviors [37]. Furthermore, glucocorticoids administered to explanted E15 mouse SCN led to earlier onset of circadian rhythms [28]. We conclude that single injections of glucocorticoids in late gestation can shift fetal circadian rhythms and daily administration can entrain fetal circadian rhythms. These data suggest future studies should test the benefits of dexamethasone delivered to women at risk of preterm birth around the time of waking when cortisol typically peaks.

We found evidence for coordinated circadian rhythms between different reproductive and fetal tissues. Previous studies have shown that the uterus and placenta have circadian rhythms during pregnancy but did not examine coordination between them [28,29,31,46]. We found that daily rhythms in the uterus consistently peak a few hours before the cervix and placenta in vitro, but around E15 these relationships become unreliable within and between pregnancies. This suggests a reorganization of the maternal and fetal circadian systems around the time of maternal-fetal circadian synchronization recorded *in vivo*. Prior studies reported circadian rhythms in maternal placenta (decidua), but conflicting results in fetal placenta layers (junctional and labyrinth zones) [28,29,49]. We found that all three placental layers have intrinsic PER2 rhythms at E15 and E18.We found highly synchronized daily rhythms between maternal and fetal layers of the E12 placenta that were sensitive to glucocorticoid agonist or antagonists in vitro, consistent with a prior observation that glucocorticoids altered placental circadian rhythms [28]. This suggests glucocorticoids synchronize daily rhythms within the placenta at least at E12. However, mifepristone also blocks progesterone signaling so we cannot rule out that progesterone may also serve an important role in synchronizing placental rhythms [50]. Further studies should provide mechanistic insights into the cells and signals mediating synchronization of maternal and fetal daily rhythms during pregnancy.

We conclude that fetal circadian rhythms appear by E9 and synchronize to the mother over the course of murine pregnancy. Synchronization to the mother and local light cycle completes around E15.5 and depends on glucocorticoids. Our findings may help explain some of the reproductive complications associated with shift work and disrupting daily rhythms during pregnancy.

## Supporting Information

**Supplementary Figure 1.**
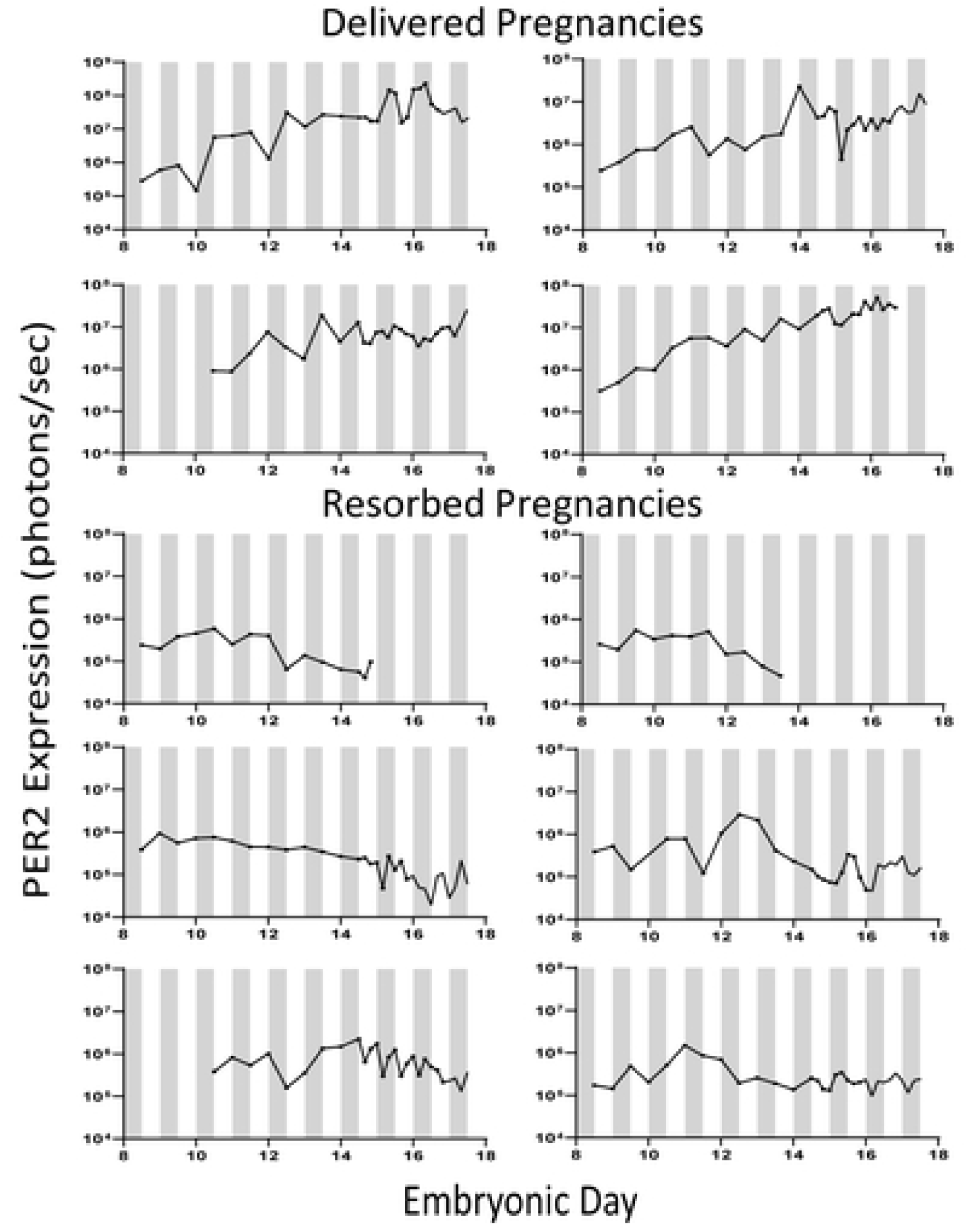
Fetal PER2 expression traces from dams that delivered (n=4) and dams that did not (n=6) following luciferin injections and bioluminescence imaging starting before E11.

**Supp. Fig. 2:**
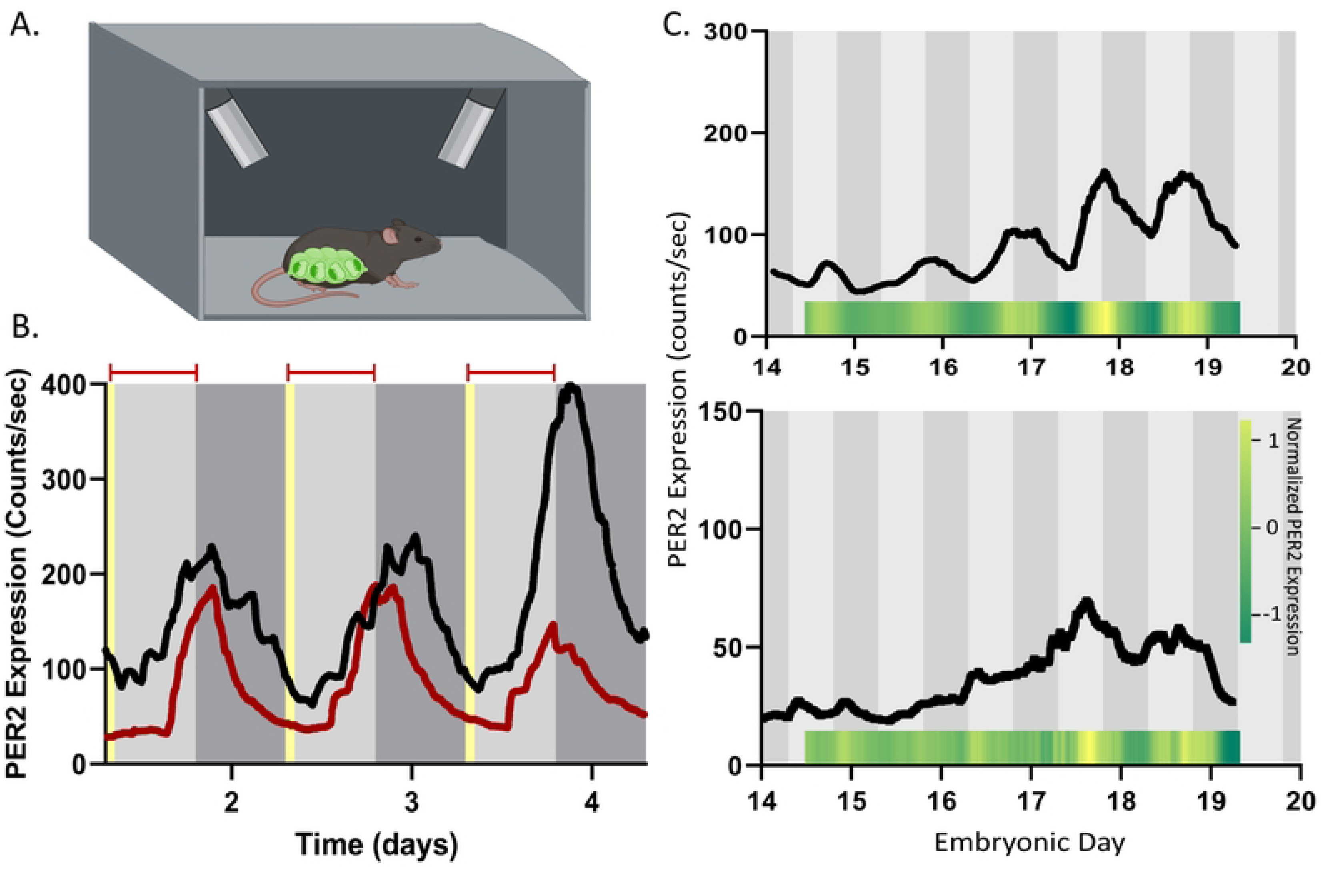
Daily rhythms in PER2 expression recorded in freely moving mice. A) Graphic of a freely moving pregnant mouse housed in a light-tight box with 2 PMTs recording fetal PER2. B. Bioluminescence from a non-pregnant, homozygous PER2::LUC mouse that received luciferin-supplemented water during the day (red line) or ad lib (black line). Bioluminescence was recorded every second except when lights were on from 7:00-7:15 am (yellow bar) each day. We detected peak PER2 expression during the early subjective night regardless of when luciferin was available. C) *In utero* fetal circadian rhythms synchronized to the mother prior to birth. Fetal PER2 from two dams (black lines and raster plots) was circadian and peaked reliably around subjective dusk by E17.

**Supp. Fig. 3.**
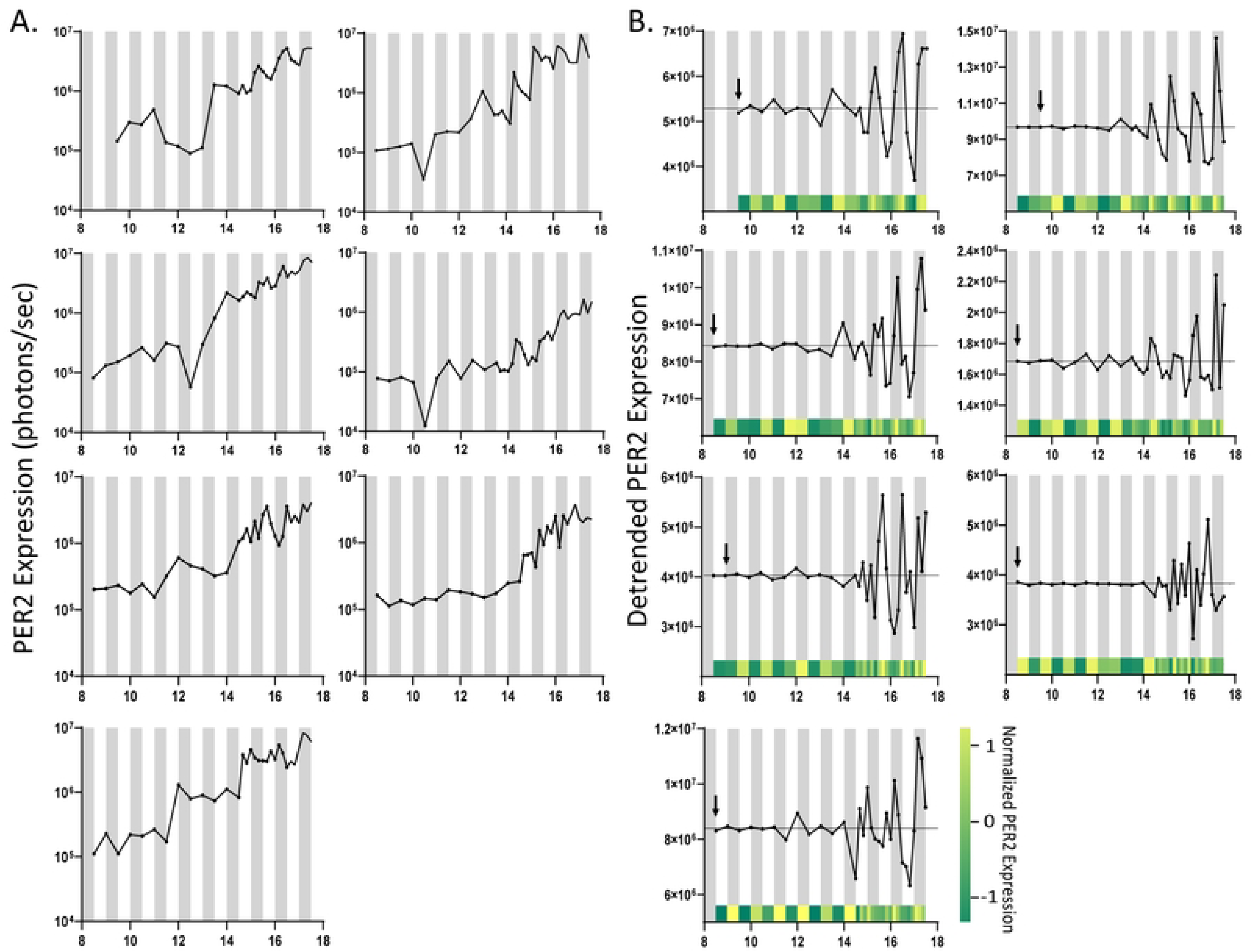
Day-night fluctuations of fetal PER2 expression. A) Fetal PER2 bioluminescence from 7 pregnancies showed exponential increases throughout gestation and variable times of peak daily expression until around E15.5. B) After subtracting the exponential increase in bioluminescence, detrended traces highlight that day-night differences in expression began by E8.5-E9.5. The arrow marks the first point when bioluminescence crossed the mean expression twice per day. The raster plot at the bottom shows PER2 expression normalized to the daily maximum to illustrate day-night differences in expression. Note the high amplitude rhythms and synchronous times of peak PER2 starting around E15.5.

**Supp. Fig. 4:**
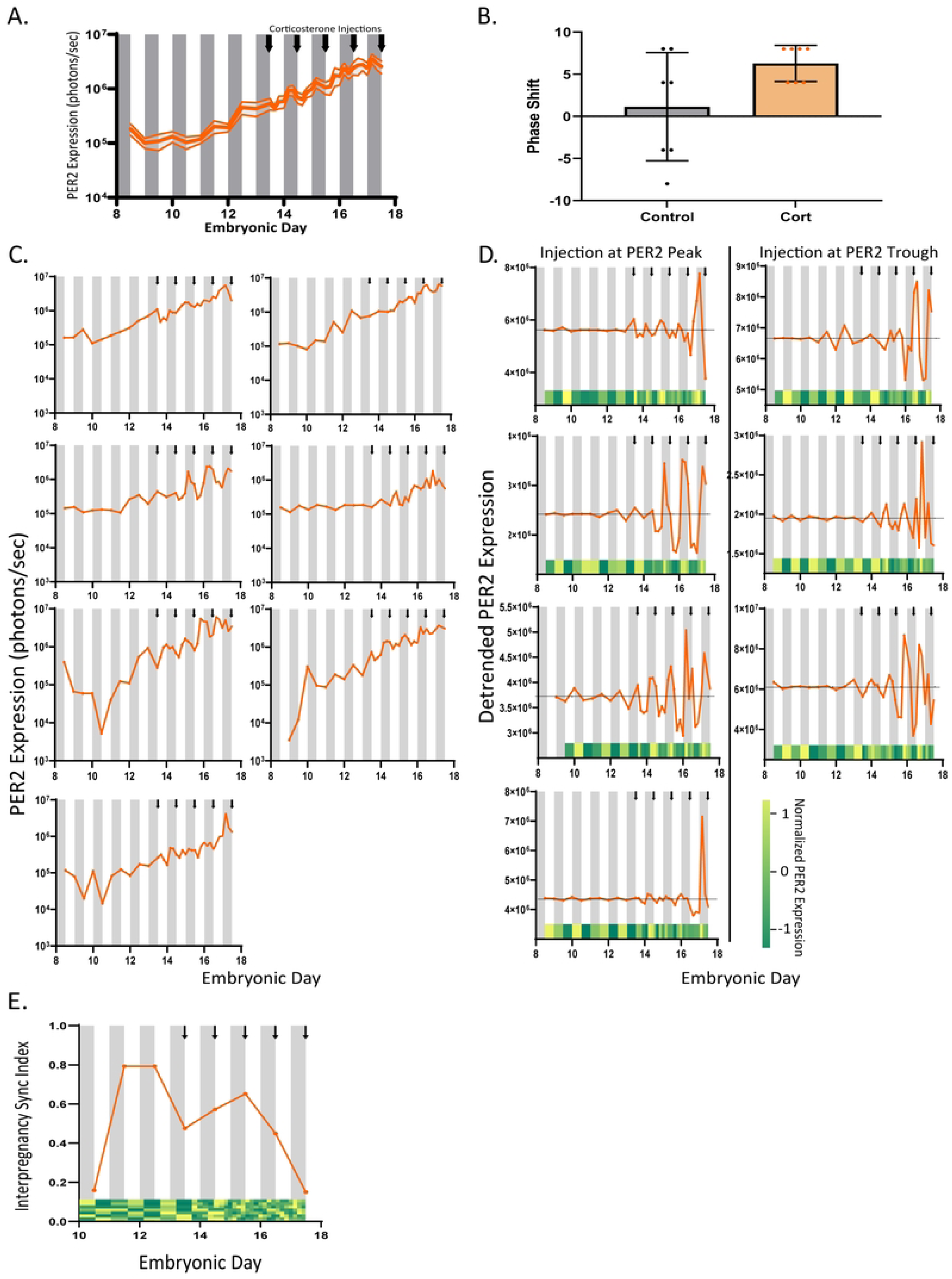
Corticosterone injections shifted fetal PER2 expression depending on the initial phase of fetal PER2. A) Average PER2 expression from pregnancies imaged from E8.5 to E17.5 (n=7). Arrows indicate when ZT0 corticosterone injections were administered (E13.5-E17.5). Corticosterone injections disrupted the average day-night differences in fetal PER2 across pregnancies. B) Daily corticosterone advanced peak fetal PER2 expression by 4-8 h compared to controls on E17.5. C) *In utero* PER2 expression from 7 corticosterone-treated pregnancies measured between E8.5 to E17.5. Arrows indicate time of corticosterone injections. D) After subtracting the 24-h average, fetal bioluminescence rhythms reveal initial corticosterone injections occurred near the trough (right) or peak (left) of fetal PER2 expression. Raster plots show the normalized detrended data. E) Interpregnancy synchrony index and raster plots illustrate how corticosterone injections disrupted synchrony between pregnancies.

**Supp. Fig. 5.**
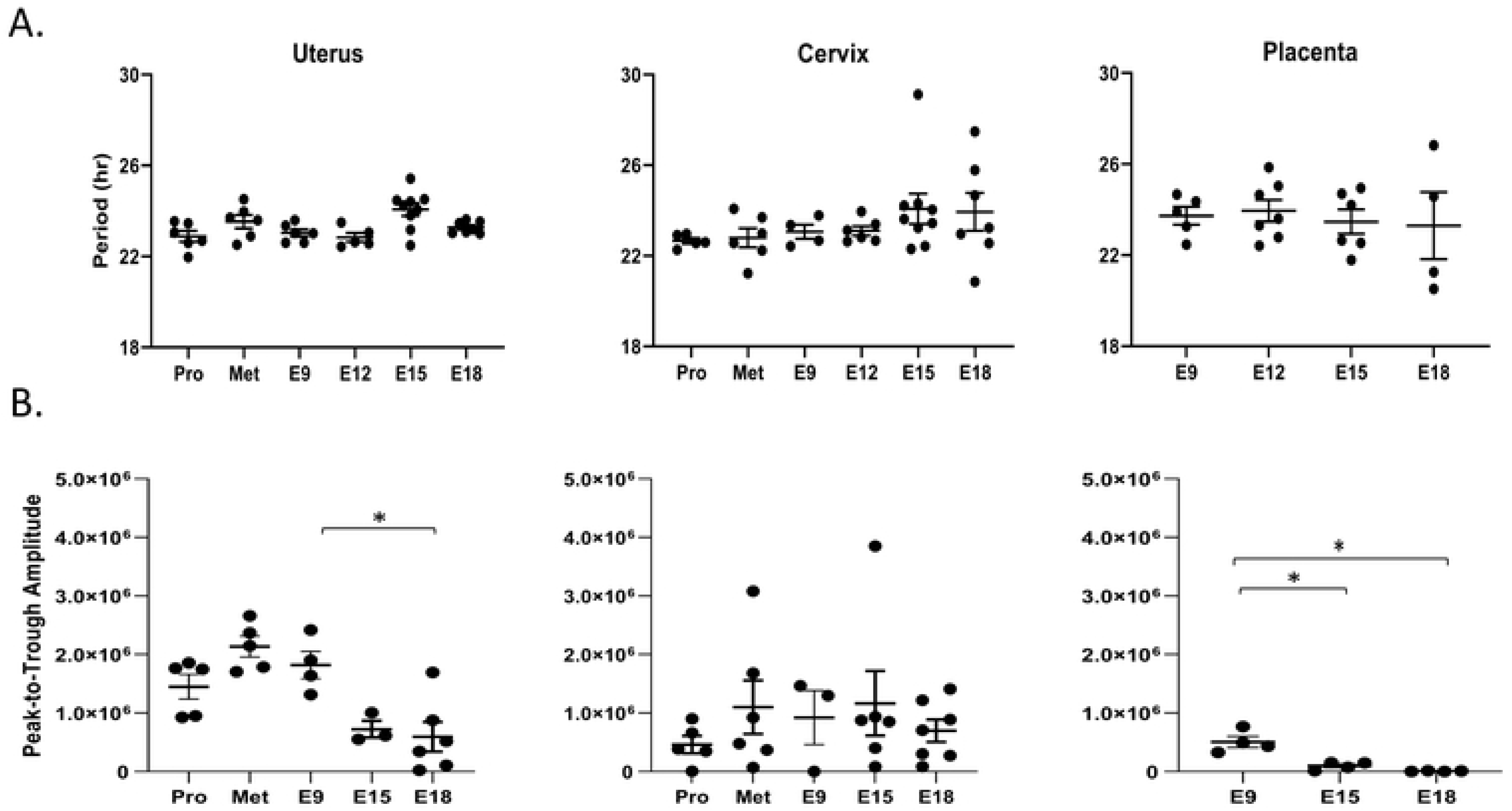
Isolated maternal and placental tissues show stable circadian periods but decreasing amplitude with gestational age. A) Circadian period did not change for uterus, cervix or placenta recorded at proestrus, metestrus, E9, E15 or E18. B) The peak-to-trough amplitude decreased in the uterus explanted on E18 compared to E9 and in the placenta explanted on E15 and E18 compared to E9 (One-way ANOVA; mean ± S.E.M).

**Supp. Fig. 6.**
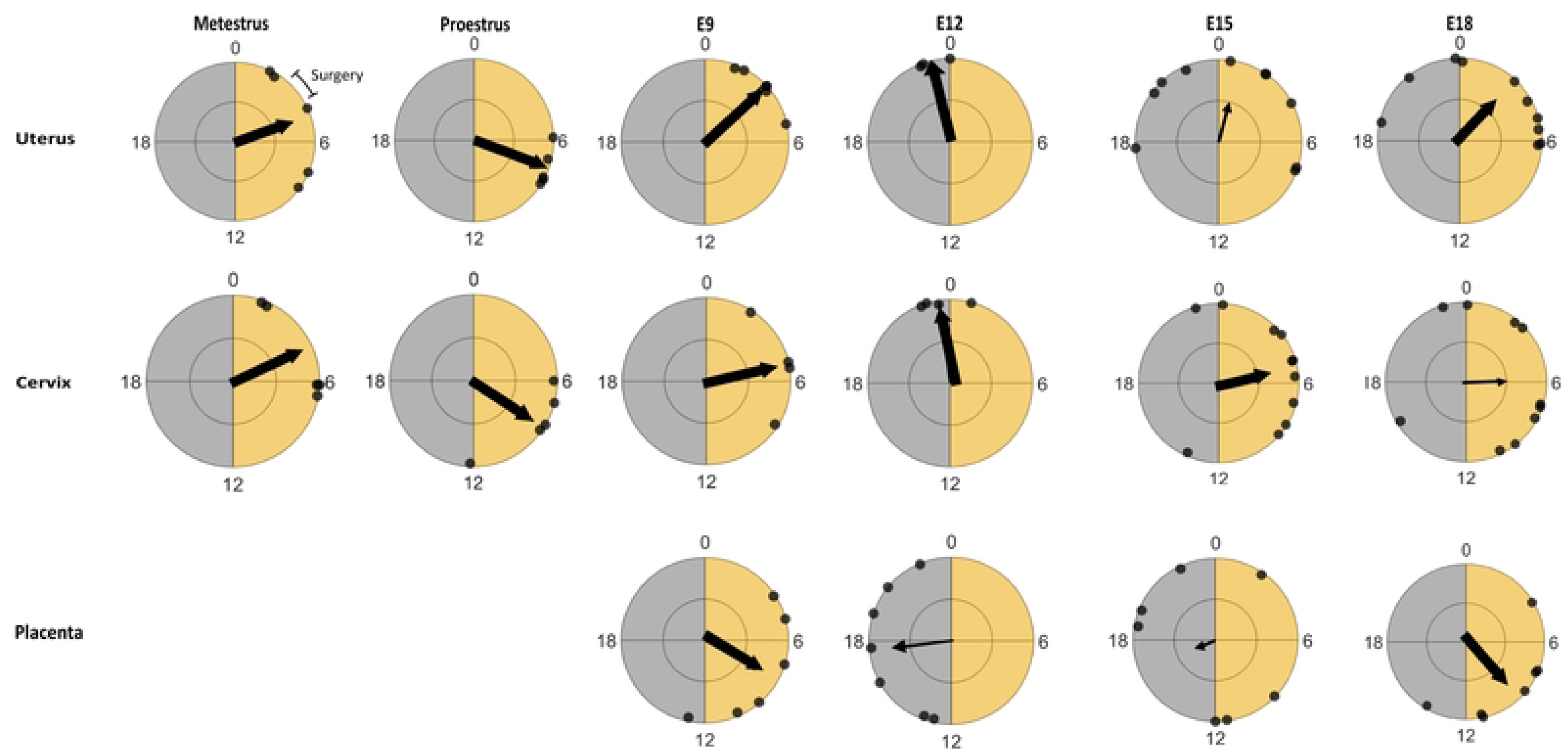
Circadian rhythms peak at reliable times in reproductive tissues depending on gestational age. Rayleigh plots reveal the average time of day when PER2 peaked (arrow) in different tissues on the second day *in vitro*. Explants (black dot) from the uterus collected on metestrus, for example, expressed average peak PER2 around ZT4, 4 h after onset of light (yellow). Large arrows show samples that had times of peak expression that significantly differed from random (i.e., significant phase clustering). PER2 expression peaked at reliable times in the uterus at metestrus (p= 0.04), proestrus (p= 0.0002), E9 (p< 0.001), E12 (p= 0.038), and E18 (p= 0.003), but not E15 (p= 0.102). In the placenta, PER2 expression reliably peaked at E9 (p= 0.047) and E18 (p= 0.004), but not E12 (p= 0.10) and E18 (p= 0.874). PER2 expression in the cervix reliably peaked at metestrus (p= 0.04), proestrus (p= 0.016), E9 (p= 0.035). E12 (p= 0.011), and E15 (p= 0.016) but not E18 (p= 0.48).

**Supp. Fig. 7.**
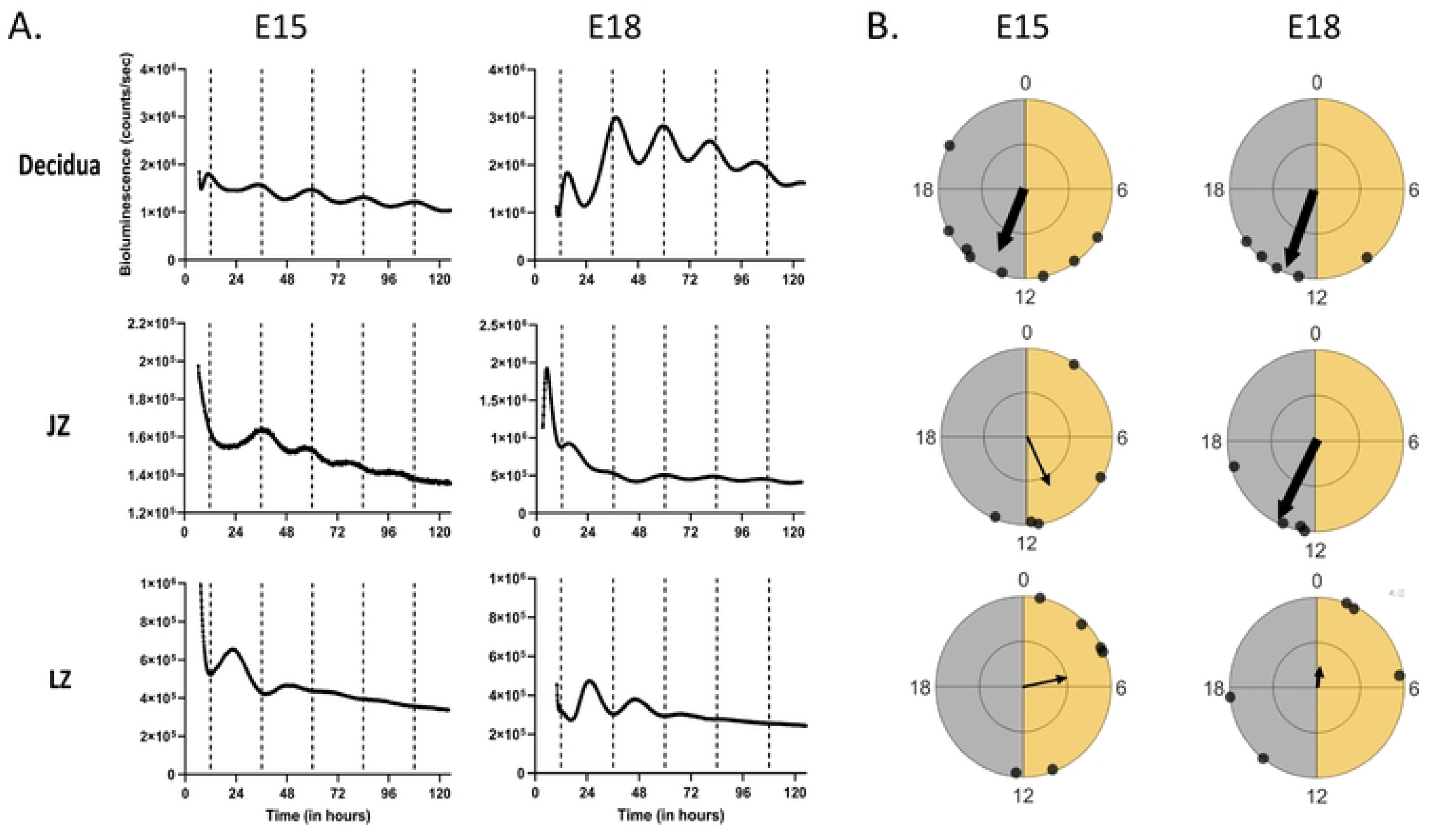
Circadian rhythms in isolated placental layers. A) PER2 bioluminescence from representative explants of the decidua, junctional zone (JZ), and labyrinth zone (LZ) collected at E15 or E18. All three layers exhibited intrinsic circadian rhythms. B) Rayleigh plots show that PER2 in the decidua peaked, on average, in the early subjective night while the other two layers were less reliable in their time of peak PER2 expression.

